# Isogenic iPSC-derived proximal and distal lung-on-chip models: Tissue- and virus-specific immune responses in human lungs

**DOI:** 10.1101/2023.11.24.568532

**Authors:** Sachin Yadav, Kazuya Fujimoto, Toru Takenaga, Senye Takahashi, Yukiko Muramoto, Ryuta Mikawa, Takeshi Noda, Shimpei Gotoh, Ryuji Yokokawa

**Author notes:** Corresponding author (R.Y).

## Abstract

Micro-physiological systems (MPS) are set to play a vital role in preclinical studies, particularly in the context of future viral pandemics. Nonetheless, the development of MPS is often impeded by the scarcity of reliable cell sources, especially when seeking various organs or tissues from a single patient for comparative analysis of the host immune response. Herein, we developed human airway-on-chip and alveolus-on-chip models using induced pluripotent stem cell (iPSC)-derived isogenic lung progenitor cells. Both models demonstrated the replication of two different respiratory viruses, namely SARS-CoV-2 and Influenza, as well as related cellular damage and innate immune responses-on-chip. Our findings reveal distinct immune responses to SARS-CoV-2 in the proximal and distal lung-on-chip models. The airway chips exhibited a robust interferon (IFN)-dependent immune response, whereas the alveolus chips exhibited dysregulated IFN activation but a significantly upregulated chemokine pathway. In contrast, Influenza virus infection induced a more pronounced immune response and cellular damage in both chip models compared to SARS-CoV-2. Thus, iPSC-derived lung-on-chip models may aid in quickly gaining insights into viral pathology and screening potential drugs for future pandemics.

---

Human lungs are prone to infections caused by respiratory viruses, including SARS-CoV-2 and Influenza A virus (IAV)^1,2^. These respiratory infections have been responsible for numerous pandemics and have placed a substantial burden on healthcare systems and management, as exemplified by the COVID-19 pandemic^3–7^. These infections affect various regions of the human lungs, from the proximal airway tract to the distal alveolar sac. Typically, a virus gains entry through cells that express virus receptors, such as multiciliated cells in the lung airway. Eventually, it reaches alveolar type I (AT1) or alveolar type 2 (AT2) cells in the alveoli, leading to alveolar diffusion and other pathological effects^8–12^. In severe cases, the infection can also spread to other organs, resulting in multiorgan damage^13–16^.

Host cells detect pathogens through “pattern recognition receptors” which, upon activation, trigger innate immune responses and generate multiple interferons (IFN) and proinflammatory cytokines^17–19^. Additionally, chemokines, which are chemotactic cytokines controlling the migration and positioning of immune cells in tissues, play a critical role in the function of the innate immune system^20,21^.

Viruses such as SARS-CoV-2 and IAV have developed strategies to evade the innate immune response, for example, by inhibiting the type I IFN pathway^22,23^. An attenuated type I IFN response can result in uncontrolled viral replication and the accumulation of immune cells at the infection site, ultimately leading to an excessive release of IFNs and the onset of a cytokine storm^24^.

Recent research findings also indicate distinct tissue-specific innate immune responses in the upper and lower respiratory tracts, especially in the context of coronaviruses such as SARS-CoV-2. These findings suggest a role for the upper airway in active viral transmission and immune defense. The diminished innate immune response in the distal lung may potentially underlie the unique uncontrolled late-phase lung damage observed in advanced COVID-19^25^.

The complications associated with the host immune response, especially following viral infections, highlight the need for urgently developing human-relevant models that can closely mimic the human lung microenvironment. Such models provide a robust platform for comprehending infection mechanisms and drug screening. While traditionally employed animal models have been widely used for drug screening, they often encounter ethical and technical challenges, including genetic differences^26^.

*In vitro* models, such as dish and insert cultures fail to accurately replicate the *in vivo* interface of human lungs. Since the onset of the recent COVID-19 pandemic, researchers have increasingly turned to micro-physiological systems (MPS) to emulate viral infections and the associated damage^27–29^. Si et al. have demonstrated the potential of MPS for drug screening and repurposing^30^. Nevertheless, these models exhibit significant irregularities, such as inadequate viral replication, atypical immune responses, and an inability to depict the suppression of immune responses by viruses such as SARS-CoV-2. Overall, these models display variability among cell types and struggle to accurately reproduce the diverse immune signatures of viral infections.

One of the primary reasons for these irregularities is the lack of a suitable cell source, which is a common limitation of lung MPS. Immortalized cells lack similarity to *in vivo* phenotypes and genotypes, rendering them unsuitable for emulating the human lung tissue interface^31^. In contrast, primary cells have limited proliferative capacity and instability under *in vitro* conditions; for example, AT2 cells tend to differentiate into AT1-like cells when cultured *in vitro*^32^. These limitations result in irregular viral replication and immune responses under *in vitro* conditions, including MPS^27,28^. Moreover, SARS-CoV-2 exhibits varying infectivity in different regions of the human lung owing to the spatial arrangement of receptor proteins^33^ and evokes different immune responses^25,34^. This underscores the critical need for a system capable of emulating different regions of the human lung to compare host immune responses and screen relevant drugs.

The integration of induced pluripotent stem cells (iPSCs) with MPS effectively overcomes the limitations of traditional cell sources and *in vitro* models, such as their failure to mimic the *in vivo* interface and their shortcomings in terms of cell availability and stability. The utilization of iPSCs offers additional advantages, including the ability to develop different regions of the human lung from a single-cell source and the potential for personalized models. In this study, we developed airway-on-chip and alveolus-on-chip models using iPSC-derived isogenic lung progenitor cells to replicate viral infections and the associated innate immune responses on-chip.

## Results

### Differentiation of iPSCs into the airway and alveolar cells on-chip

We utilized iPSC-derived isogenic carboxypeptidase M (CPM+) lung progenitor cells to establish airway-on-chip and alveolus-on-chip models (Fig. 1a). Briefly, iPSCs underwent a 21-day differentiation process on a culture plate, and magnetic-activated cell sorting (MACS) was employed to isolate CPM+ lung progenitor cells. These CPM+ cells were then seeded into the airway and alveolus chips with the respective culture medium (Fig. S1).

**Figure 1:**
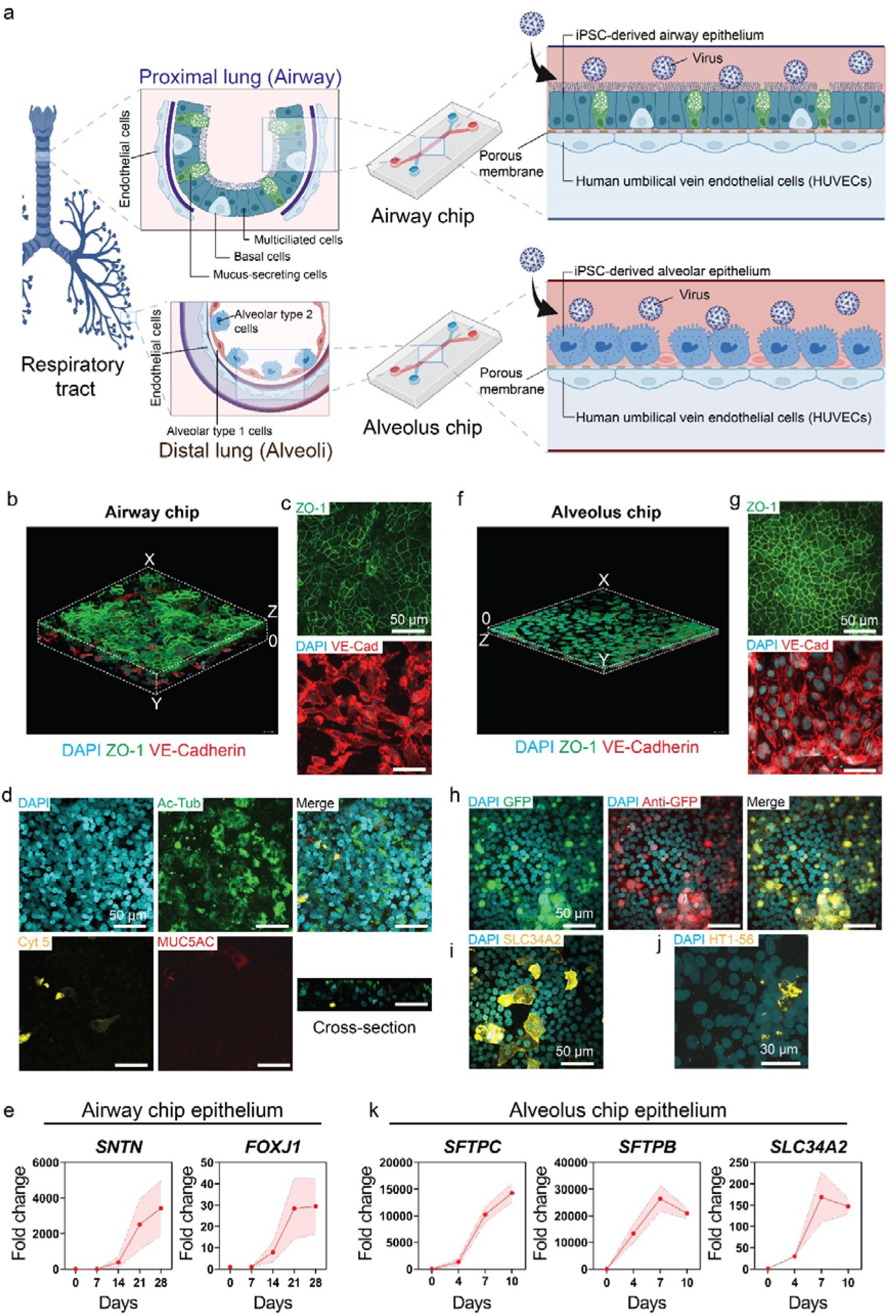
Development of human iPSC-derived airway chips and alveolus chips. Human iPSC-derived CPM+ lung progenitor cells were seeded in the top channel for further differentiation into airway and alveolar cells in the respective chips before seeding the HUVECs on the opposite side of the porous membrane. (a) Schematic illustration of the airway and alveolus chips (created with BioRender.com). (b) 3D volume view of tight junctions immunoassayed for ZO-1 and VE-cadherin in the airway epithelium and endothelium, respectively. (c) Maximum intensity z-stack immunofluorescent staining for ZO-1 and VE-Cadherin in airway epithelium and endothelium respectively. (d) Immunofluorescent staining for Ac-Tub, MUC5AC, Cyt 5 in airway epithelium. (e) Gene expression levels of *SNTN* and *FOXJ1* in the airway epithelium (n = 3, except day 7, n = 2). (f) 3D volume view of tight junctions immunoassayed for ZO-1 and VE-cadherin in the alveolus epithelium and endothelium, respectively. (g) Maximum intensity z-stack immunofluorescent staining for ZO-1 and VE-Cadherin in alveolus epithelium and endothelium respectively. (h) GFP expression and Immunofluorescence staining of GFP in the alveolus epithelium. (i) Immunofluorescence staining of SLC34A2 in alveolus epithelium. (j) Immunofluorescence staining of HT1-56 in alveolus epithelium. (k) Gene expression levels of *SFTPC*, *SFTPB*, *SLC34A2* in the alveolus epithelium (n = 3). (Immunofluorescence images were taken on day 32 for airway chips and day 7-9 for alveolus chips).

Immunofluorescence staining of ZO-1 and VE-Cadherin revealed the formation of tight junctions in both the epithelium and endothelium tissues of the airway chip respectively (Fig. 1b, c). The optimal differentiation of (CPM+) cells into multiciliated cells was confirmed by immunofluorescent staining of the airway multiciliated cell marker acetylated tubulin (Ac-Tub). Additionally, other airway epithelial cells such as mucus-secreting cells (MUC5AC) and basal cells (Cytokeratin 5) were also observed (Fig. 1d), consistent with prior reports^35^. Gene expression analysis of key multiciliated cell markers, including *SNTN* and *FOXJ1*, indicated progressive differentiation of CPM+ lung progenitor cells into multiciliated cells by day 28 (Fig. 1e). Furthermore, the presence of mucus-secreting club cells was verified by the upregulation of *SCGB1A1 (CCSP)* and *SPDEF* genes (Fig. S2a).

Similarly, immunofluorescent staining of ZO-1 and VE-Cadherin showed the formation of tight junctions in both the epithelium and endothelium tissues of the alveolus chip respectively (Fig. 1f, g). Given that an SFTPC-GFP knock-in iPSC line^36^ was utilized in the experiments, alveolar type 2 (AT2) cells expressed GFP and were detected using an anti-GFP antibody (Fig. 1h). The differentiation of CPM+ cells into AT2 cells was further confirmed through immunofluorescent staining of SLC34A2 (Fig. 1i), a key marker for AT2 cells. Additionally, some alveolar type 1 (AT1)-like cells were observed in the alveolar epithelium by staining HT1-56 (Fig. 1j). Gene expression analysis of key AT2 cell markers, including *SFTPC, SFTPB,* and *SLC34A2*, demonstrated the progressive differentiation of CPM+ cells into AT2 cells by day 7 (Fig. 1k). Gene expression of AT1 cell marker (*AGER* and *PDPN*) also increased from day zero to day 10 (Fig. S2b). Given that multiciliated cells and AT2 cells are the primary targets of SARS-CoV-2 in the proximal airway and alveoli, respectively, our airway chip and alveolus chip offer an excellent means to replicate viral pathogenesis and host immune responses.

### Both chips exhibit high susceptibility to SARS-CoV-2 with minimal cellular damage

Transmembrane proteins, namely angiotensin-converting enzyme 2 (ACE2), transmembrane serine protease 2 (TMPRSS-2), and neuropilin-1 (NRP-1), play a pivotal role in facilitating the entry of SARS-CoV-2 into human host cells^37,38^. The gene expression of all three genes, *ACE2*, *TMPRSS-2*, and *NRP-1* increased progressively as CPM+ lung progenitor cells differentiated from day zero to day 28 in airway epithelium and from day zero to day 10 in alveolus epithelium (Fig. 2a, b).

**Figure 2:**
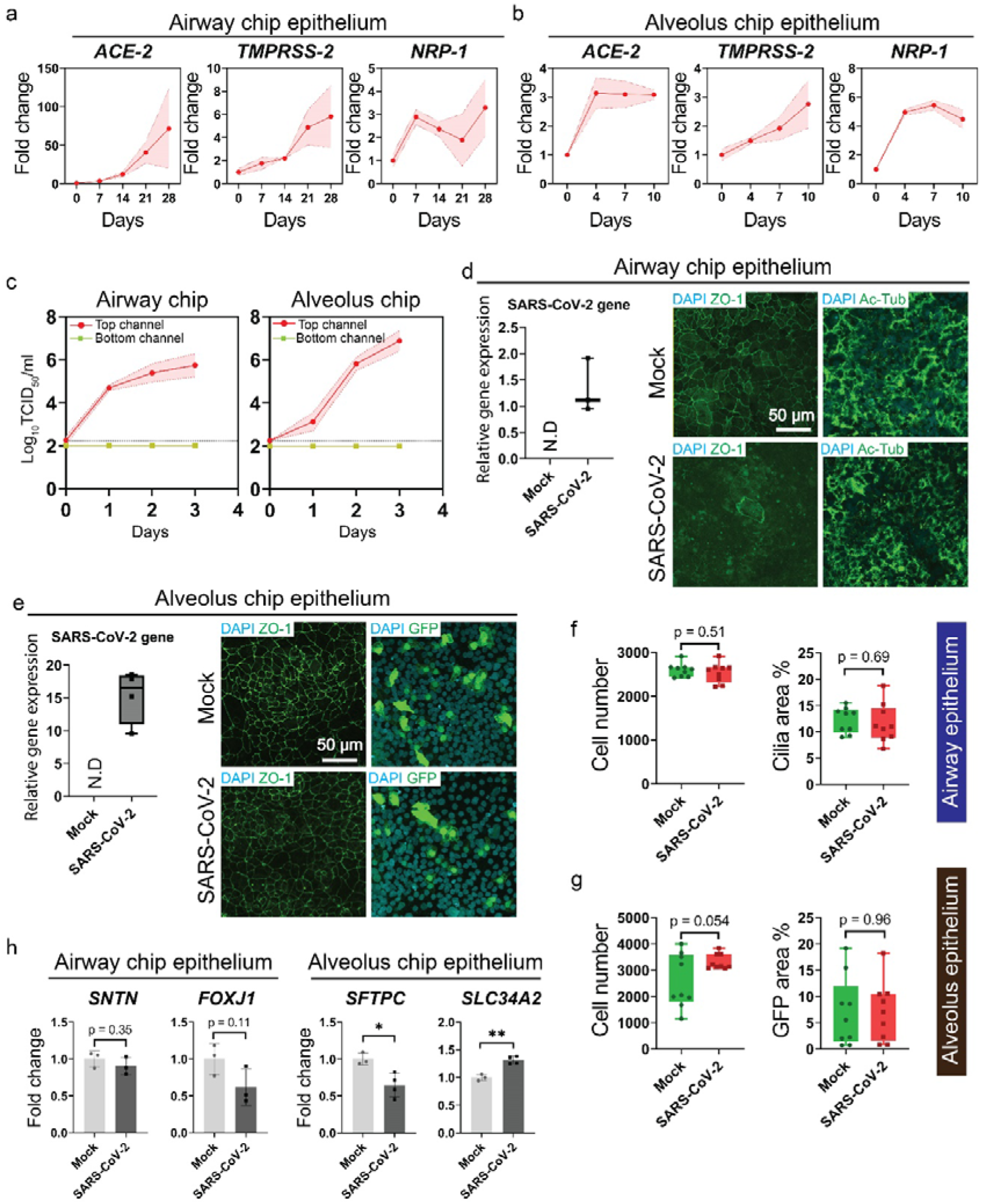
SARS-CoV-2 infection of airway chips and alveolus chips. SARS-CoV-2 was introduced into the top channel of both chips at a concentration of 10^3^ TCID_50_/device, and the devices were cultured for three days. (a) Gene expression of the SARS-CoV-2 receptors *ACE-2*, *TMPRSS-2*, and *NRP-1* in the airway epithelium measured on day 0, 7, 21, and 28 (n = 3 chips, except day 7, n = 2). (b) Gene expression of the SARS-CoV-2 receptors *ACE-2*, *TMPRSS-2*, and *NRP-1* in the alveolus epithelium measured on day 0, 4, 7, and 10 (n = 3 chips except *NRP-1*, day 0 = 2). (c) Viral titers in the airway and alveolus chips were measured at 3dpi using a 50% tissue culture infectious dose (TCID_50_) method (n of at least 3 chips). (d) Intracellular SARS-CoV-2 gene expression in airway chip epithelial cells normalized to housekeeping gene *ACTB* and immunofluorescent staining for ZO-1, and Ac-Tub in mock and infected airway epithelium at 3 dpi. (e) Intracellular SARS-CoV-2 gene expression in alveolus chip epithelial cells normalized to housekeeping gene *ACTB* and immunofluorescent staining for ZO-1, and GFP in mock and infected alveolus epithelium at 3 dpi. (f) Image analysis of immunofluorescence staining in mock and infected airway chip samples to determine the cell number and cilia area percentage (Ac-Tub) (n = at least three ROI from independent 3 chips). (g) Image analysis of immunofluorescence staining in mock-and infected alveolus chip samples for cell number and GFP area percentage (n = at least three ROI from independent 3 chips). (h) Gene expression of key cellular markers *SNTN* and *FOXJ1* in the airway epithelium and *SFTPC* and *SLC34A2* in the alveolus epithelium in mock and infected conditions at 3 dpi (n = 3).

SARS-CoV-2 was introduced on day 32 in the top channel of the airway chips and on day 8 in the alveolus chips, respectively. The viral titer in the epithelium channel of the respective chips increased from day 0 to day 3 post-infection, while no viral replication was detected in the endothelial channel, consistent with previous reports^39,40^ (Fig. 2c). The presence of intracellular SARS-CoV-2 viral transcripts was also observed at 3dpi (Fig. 2d). SARS-CoV-2 infection resulted in the disruption of tight junctions between infected cells with minimal decrease in number of ciliated cells (Fig. 2d).

Similarly, SARS-CoV-2 infection in the alveolus epithelium was confirmed by the presence of intracellular SARS-CoV-2 viral transcripts at 3dpi (Fig. 2e). Interestingly, SARS-CoV-2 infection did not disrupt the tight junctions of the alveolus epithelium, and there was only a minimal decrease in the GFP expression of AT2 cells (Fig. 2e). The cell number did not change significantly after SARS-CoV-2 infection in either airway or alveolus epithelium (Fig. 2f, g). Comparatively, the cilia area percentage in mock and SARS-CoV-2 infected airway chips showed no significant difference (Fig. 2f). The GFP area percentage quantification revealed an insignificant change in the number of GFP-positive cells (Fig. 2g).

While SARS-CoV-2 infection led to a decrease in the expression of both *SNTN* and *FOXJ1*, the changes were not statistically significant (Fig. 2h). In alveolus chips, SARS-CoV-2 infection resulted in a significant decrease in the expression of *SFTPC*, whereas the expression of *SLC34A2* increased after infection (Fig. 2h).

### RNA sequencing unveils a dysregulated type I IFN activation and strong chemokine response in alveolus epithelium

In the airway epithelial cells, RNA sequencing at 3 days post-infection revealed the upregulation of 165 genes and the downregulation of 40 genes. The volcano plot depicted the activation of innate immune response-related genes, including *IRF*s, *ISG*s, and *IFN*s (Fig. 3a). Gene ontology (GO) analysis unveiled the upregulation of key biological processes related to the immune response, particularly the type I IFN pathway. Among the upregulated pathways, the response to interferon beta, interferon alpha, positive regulation of interferon beta production, and interferon-gamma (type II IFN pathway) were the most significant. Additionally, SARS-CoV-2 infection upregulated the MHC Class 1 related pathway, indicating an overall robust innate immune response in airway epithelium (Fig. 3b). A robust type I IFN activation, crucial for the host cell’s antiviral response, was indicated by the upregulation of negative regulation of viral genome replication (Fig. 3b). Further pathway analysis using Reactome indicated the activation of interferon signaling through multiple processes, confirming SARS-CoV-2 infection (Fig. S3).

**Figure 3:**
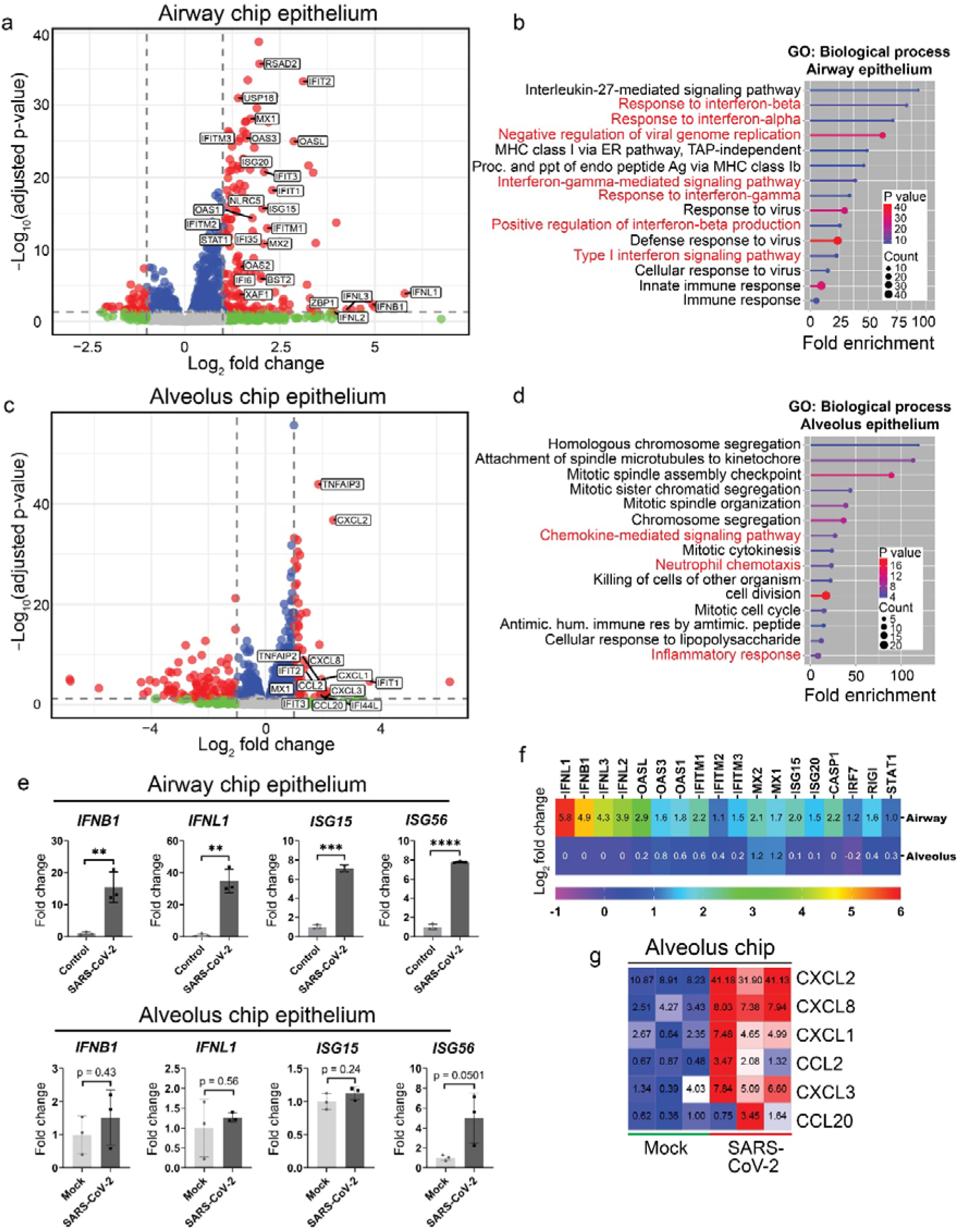
RNA sequencing reveals a dysregulated IFN activation and robust chemokine response in the alveolus epithelium. (a) Volcano plot of differentially expressed genes between mock and infected airway epithelium (log_2_ fold change > 1 and (P_adj_) < 0.05, n = 3 chips). (b) GO analysis showing the most significantly upregulated biological pathways in airway chips (P value shows −log_10_(P value). (c) Volcano plot of differentially expressed genes between mock and infected alveolus epithelium (log_2_ fold change > 1 and (P_adj_) < 0.05, n = 3 chips). (d) GO analysis showing the most significantly upregulated biological process pathways in the alveolus chips (P value shows −log_10_(P value). (e) Expression of IFN-related genes in the airway alveolus epithelium in mock and infection conditions at 3 dpi (n = 3). (f) Log_2_ fold change heatmap showing the expression of important innate immune response genes in the airway and alveolus epithelium of mock and infected samples (n = 3). (g) Transcripts per million (TPM) heatmap of chemokine gene expression in mock and infected alveolus epithelium on alveolus chips.

In the alveolus epithelium, RNA sequencing following SARS-CoV-2 infection resulted in the upregulation of 65 genes and the downregulation of 98 genes. The volcano plot indicated limited upregulation of innate immune response genes compared to the airway epithelium (Fig. 3c). GO analysis revealed limited activation of the immune response, particularly type I IFN pathways (Fig. 3d). Analysis of biological processes unveiled a significant activation of pathways related to immune cell recruitment, such as neutrophil chemotaxis and chemokine-mediated signaling pathways, which played a major role in the inflammatory response elicited by alveolus epithelium (Fig. 3d). Reactome analysis did not show a significant increase in any immunity-related molecular pathways, further indicating limited IFN activation in alveolus epithelium (Fig. S3). The absence of an innate immune response, particularly type I IFN activation, resulted in a muted increase in antiviral activities by host cells.

SARS-CoV-2 infection significantly increased the expression of genes related to innate immune responses, including *ISG15*, *ISG56*, *IFNL1*, and *IFNB1* in airway epithelium. Conversely, alveolus epithelium did not exhibit a significant increase in the gene expressions of *ISG15, ISG56, IFNL1, and IFNB1* after SARS-CoV-2 infection, further confirming the lack of innate immune activation in alveolus chips (Fig. 3e).

The heatmap analysis revealed the upregulation of key type I IFN genes in airway chips, while no upregulation was observed in the alveolus epithelium after SARS-CoV-2 infection (Fig. 3f). Interestingly, the investigation of chemokine-related genes in the alveolus epithelium after infection showed a robust upregulation of key chemokine-related genes in alveolus chips (Fig. 3g), possibly responsible for activating chemotaxis pathways, as indicated in the GO analysis. A similar observation was previously reported in distal *in vivo* lung samples with SARS-CoV-2 infection^34^.

### IAV causes heightened cellular damage in both airway and alveolus chips

In the airway chips, the titer of IAV increased from 1 dpi to 3 dpi in the top channel, while no viral replication was observed in the lower channel (Fig. 4a). Immunofluorescent staining of the airway chips confirmed IAV infection in epithelial cells. IAV infection resulted in significant damage to the tight junctions of the airway epithelium and a decrease in multiciliated cells (Ac-Tub) (Fig. 4b). Basal cells, which are adult stem cells located at the base of the airway epithelium near the basement membrane, can increase in number and differentiate into multiciliated cells in the event of an injury^41^. Immunofluorescent staining of Cytokeratin 5, a marker for basal cells, revealed an increase in the number of basal cells after infection (Fig. 4b). We also found a co-localization of IAV nucleoprotein antibody with Cytokeratin-5. Additionally, IAV infection caused substantial damage to the stratified layer in the airway chips, exposing the basal cells to the apical side of the cell layer (Fig. 4c). Quantitative analysis of the immunostaining images confirmed a decrease in the cell number, layer thickness, and cilia area percentage on the apical side of the airway epithelium (Fig. 4d).

**Figure 4:**
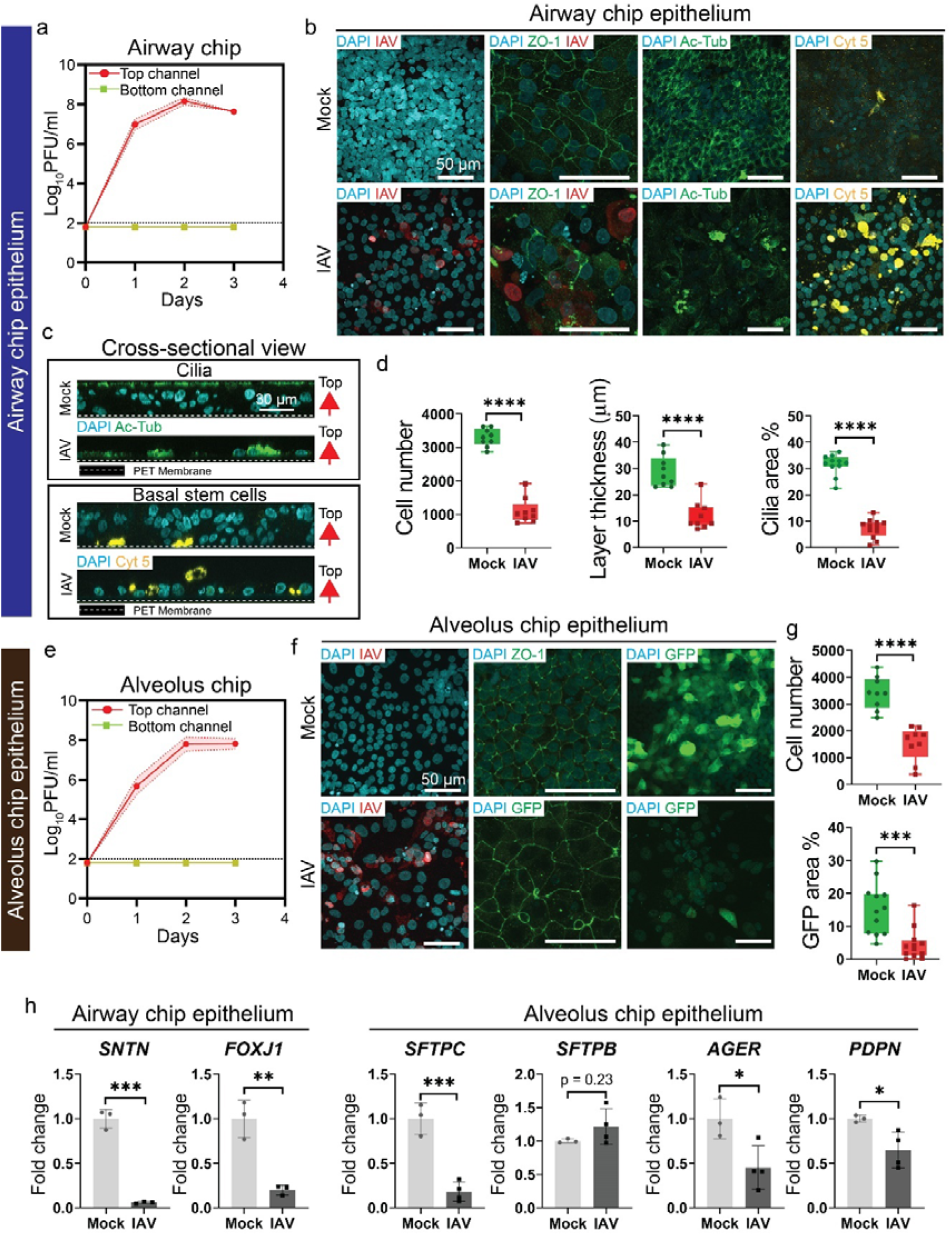
IAV inflicts multifold cellular damage and immune response in both airway and alveolus epithelium in respective chips. IAV was introduced into the top channel of both chips at a concentration of 10^3^ PFU/device, and the devices were cultured for three days. (a) The influenza viral titer in the airway chip was measured at 3 dpi using a plaque-forming assay (n = 3). (b) Immunofluorescence staining of the IAV N protein, Ac-Tub, ZO-1, and Cytokeratin 5 in mock and infected airway epithelium in airway chips. (c) Orthogonal view of immunofluorescence staining of Ac-Tub and Cytokeratin 5 in mock and infected airway epithelium in airway chips. (d) Image analysis of immunofluorescence staining of mock and infected samples to determine the cell number, layer thickness, and cilia area percentage (n = at least three ROI from three independent chips). (e) Influenza viral titer in the alveolus chip measured at 3 dpi using a plaque forming assay (n = 3). (f) Immunofluorescence staining of Influenza N protein, ZO-1, and GFP in the mock and infected alveolus epithelium of alveolus chips. (g) Image analysis of immunofluorescent staining in mock and infected samples for the number of nuclei and GFP area percentage (n = at least three ROI from independent three chips). (h) Expression of *SNTN, FOXJ1, SFTPC, SFTPB, SLC34A2, AGER,* and *PDPN* in respective chips of mock-and infected samples, (n = 3).

Like the airway chips, the titer of IAV increased from day 1 to day 3 post-infection in the top channel of the alveolus chips, with no viral replication observed in the bottom channel (Fig. 4e). Immunofluorescent staining of the alveolus chips revealed IAV infection in epithelial cells (Fig. 4f). IAV infection resulted in an increase in cell size but left tight junctions intact (Fig. 4f). IAV caused substantial damage to AT2 cells, resulting in a decrease in GFP-positive cells. Quantitative analysis of the immunostaining images confirmed a decrease in cell number and GFP area percentage in the alveolus epithelium (Fig. 4g).

Gene expression of key multiciliated cells markers, *SNTN* and *FOXJ1*, was significantly reduced after IAV infection in airway chips. While in alveolus chips, IAV infection led to a significant decrease in key markers for AT2 cells, such as *SFTPC*. AT1 cell markers *AGER* and *PDPN* were also significantly reduced after IAV infection (Fig. 4h). Interestingly, *SFTPB* gene expression did not show any significant changes, while *SLC34A2* exhibited a significant increase.

### IAV indiscriminately activates type I pathway in both airway and alveolus chips

IAV infection resulted in the upregulation of 2271 genes and the downregulation of 3125 genes in airway epithelium, while in alveolus epithelium, IAV infection upregulated 1432 genes and downregulated 930 genes (Fig. 5a, b). Notably, IFNs were upregulated in both chips. A heatmap comparison further confirmed the upregulation of crucial type I IFN genes, including *IFNL1, IFNL2, IFNB1, IFNL3, OASL, ISGs, IFITMs, RIGI, IRF7,* and *STAT1* in both airway and alveolus epithelium (Fig. 5c). Gene ontology analysis revealed the activation of pathways such as negative regulation of virus replication, defense response against the virus, and innate immune response, among others, in both chips (Fig. 5e). In line with our previous observations using immunofluorescence assay (IFA), the GO of downregulated genes indicated a decrease in cell-cell adhesion and extracellular matrix (ECM) production in alveolus chips, while significant damage was observed to multiciliated cells and the cytoskeleton in airway chips after IAV infection (Fig. 5d). These findings were further validated using RT-PCR to demonstrate the activation of *ISG15*, *IFNL1*, and *IFNB1* in both chips (Fig. 5e, f). The PCA plot displayed extreme coordinates in both airway and alveolus epithelium after IAV infection compared with SARS-CoV-2 infection (Fig. 5g).

**Figure 5:**
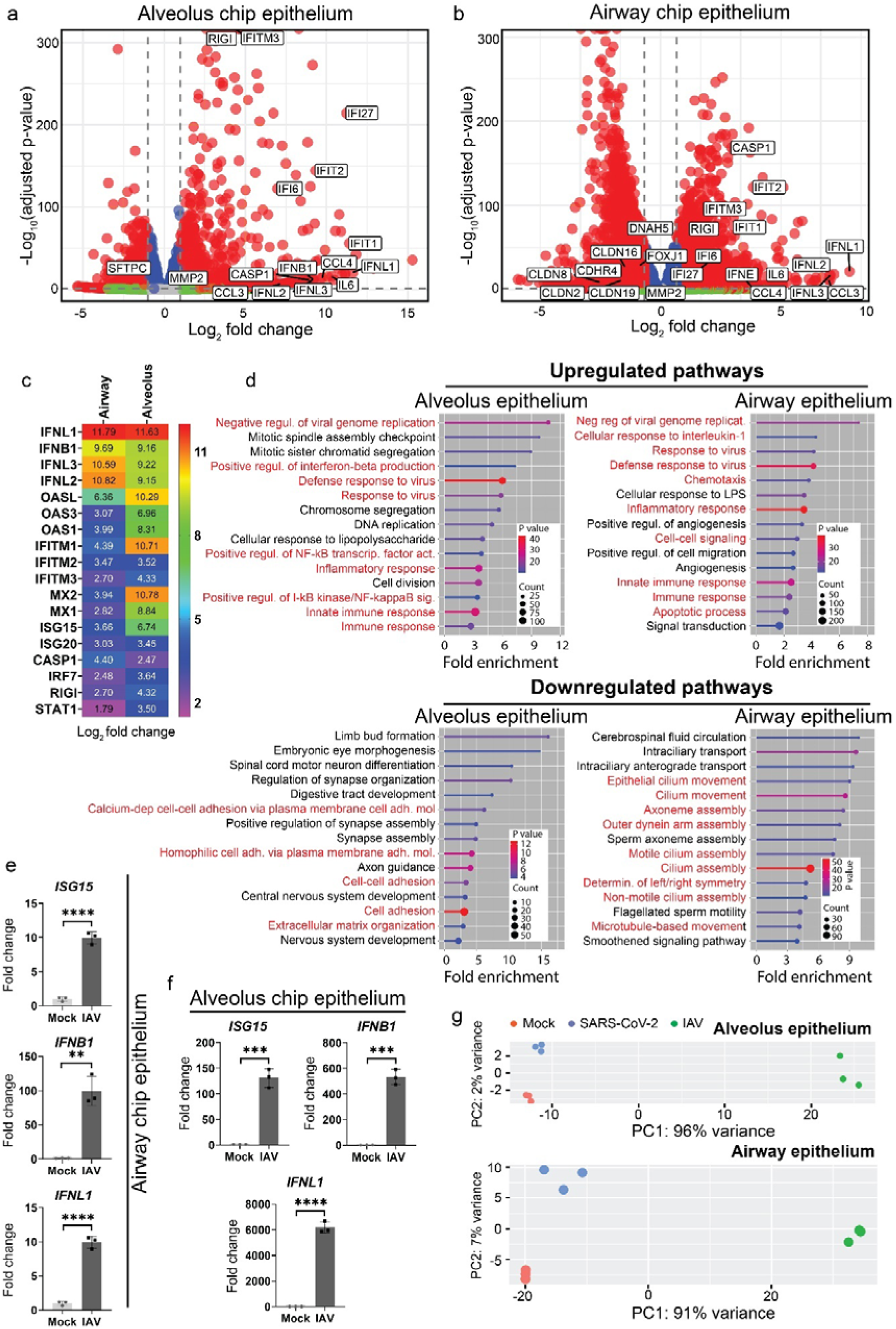
IAV elicits a robust immune response in both airway chip and alveolus chip unlike SARS-CoV-2. (a) Volcano plot of differentially expressed genes between mock and infected alveolus epithelium (log_2_ fold change > 1 and (P_adj_) < 0.05, n = 3 chips). (b) Volcano plot of differentially expressed genes between the mock and infected airway epithelium (log_2_ fold change > 1 and (P_adj_) < 0.05, n = 3 chips). (c) Log_2_ fold change heatmap showing the expression of important innate immune response genes in the airway and alveolus epithelium of mock and infected samples (n = 3). (d) GO analysis showing significantly upregulated and downregulated pathways in the respective chips between the mock and infected samples (P value shows −log_10_(P value)). (e) Expression of the key IFN genes *ISG15*, *IFNL1*, *IFNB1* in mock and infected airway epithelium in airway chips (n = 3). (f) Expression of the key IFN genes *ISG15*, *IFNL1*, and *IFNB1* in mock and infected alveolus epithelium in alveolus chips (n = 3). (g) Principal component analysis (PCA) plot of mock and infected airway and alveolus chips (n = 3).

### IAV induces an immune response in endothelial cells

Endothelial cells in both chips were not infected with SARS-CoV-2 or IAV (Fig. 2c, 3a, 3c). In previous reports, endothelial cell infection has been rarely observed^28,30,39^. Subsequently, we explored the impact of SARS-CoV-2 infection and IAV on both airway and alveolus endothelium using RNA sequencing analysis. Interestingly, no significant gene expression changes were observed in either the airway or alveolus chip endothelium after SARS-CoV-2 infection (Fig. S4a, b). However, after IAV infection, endothelial cells triggered a robust innate immune response with the upregulation of innate genes, such as *OASs, ISGs,* and *IFITMs*, in both airway and alveolus chips (Fig. 6a). GO analysis indicated the activation of type I interferon and other innate immune response pathways in the endothelium of both the airway and alveolus chips following IAV infection (Fig. S4c).

**Figure 6:**
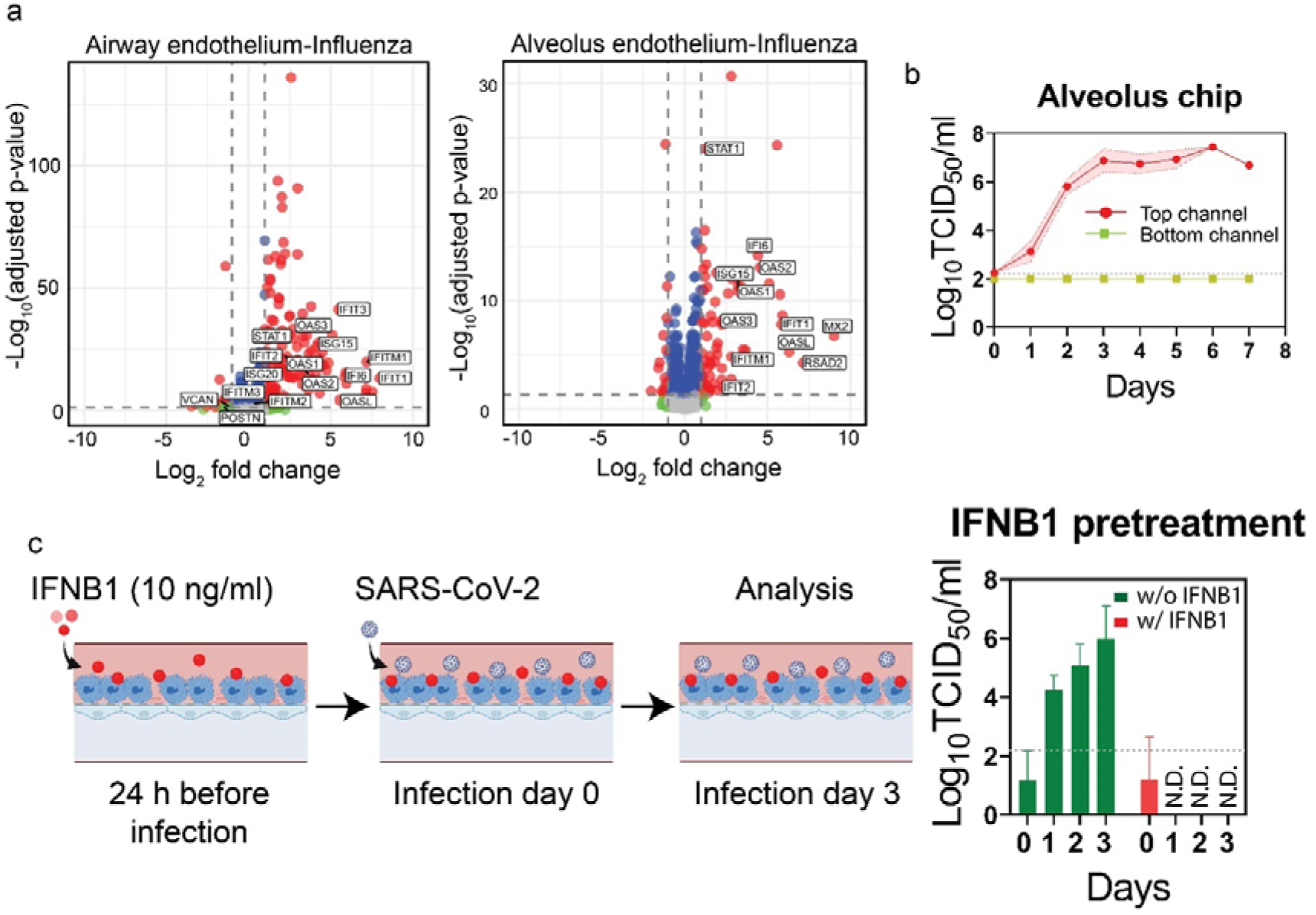
IAV induces innate immune response in endothelial cells. (a) Volcano plot of differentially expressed genes between mock and IAV-infected airway and alveolus endothelium (log_2_ fold change > 1 and (P_adj_) < 0.05, n = 3 chips). (b) SARS-CoV-2 viral titer in the alveolus chip was measured at 7 dpi using a plaque-forming assay (n = 4). (c) SARS-CoV-2 viral titer in the alveolus chips was measured with and without IFNB1 pretreatment after 3 dpi (n = 3 chips; one chip without IFNB1 pretreatment infected with SARS-CoV-2 was excluded owing failure to replicate the virus).

**Figure 7:**
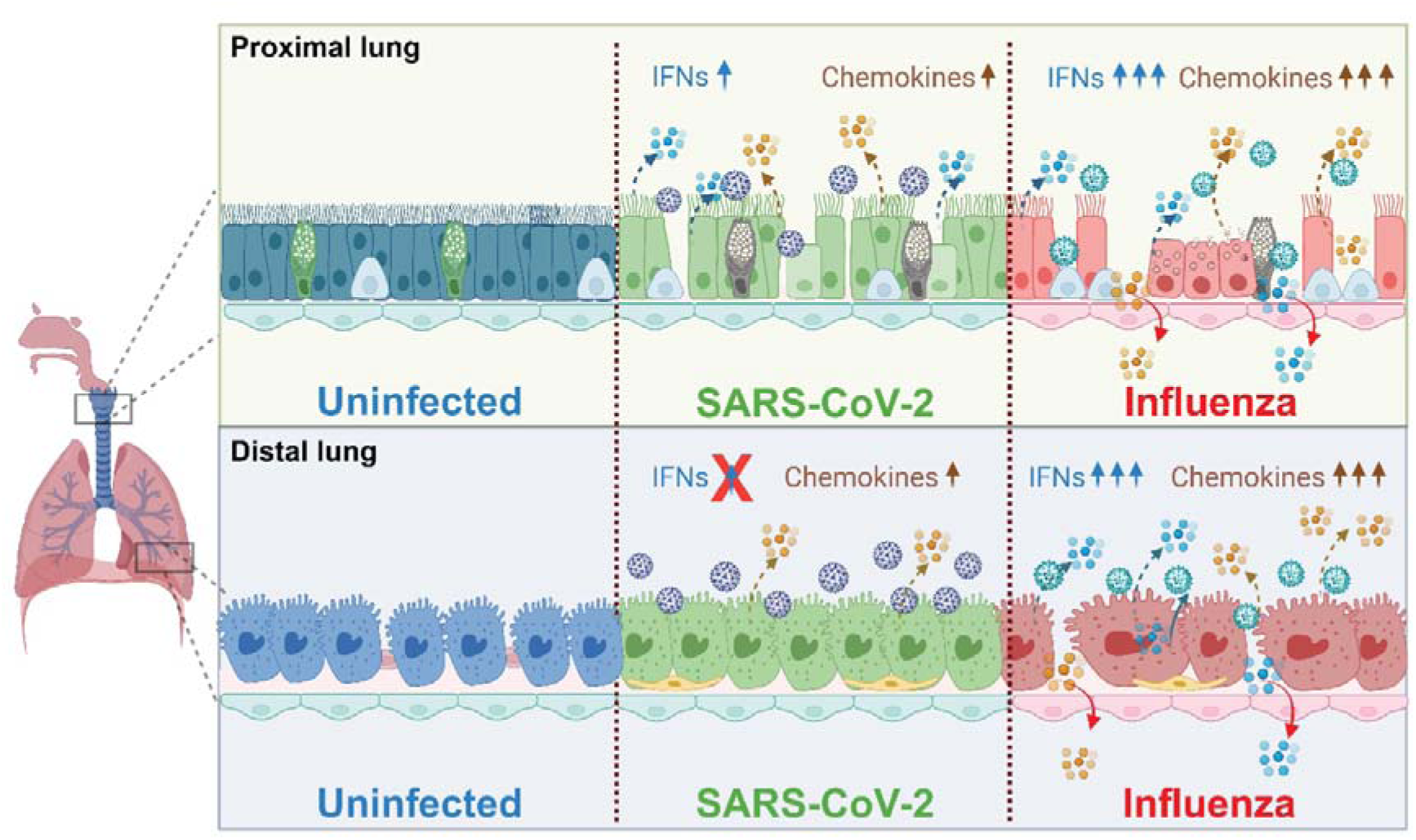
Schematic of the host immune response against SARS-CoV-2 and IAV in proximal and distal lung. In airway chips, SARS-CoV-2 infection induces an upregulation of type I IFN and chemokine pathways. Conversely, in alveolus chips, the type I IFN response is suppressed, and only the chemokine pathways are upregulated. In the case of Influenza infection, there is a significant upregulation of type I interferon pathways in both airway and alveolus chips, resulting in overwhelming cellular damage in both proximal and distal lung (Created with BioRender.com).

### IFNB1 inhibits viral replication in alveolus chips

To investigate the impact of stunted IFN activation in alveolus epithelium, SARS-CoV-2 infected chips were cultured for 7 dpi, revealing consistent viral production in the top channel (Fig. 6b). Notably, IFNB1 is considered one of the therapeutic options and has been shown to inhibit SARS-CoV-2 replication^42^. Alveolus chips pretreated with IFNB1 showed a significant inhibition of SARS-CoV-2 replication as compared to the untreated alveolus chips (Fig. 6c).

## Discussion

We developed human airway and alveolus chip models utilizing isogenic iPSCs for the first time to overcome shortcomings of immortalized and primary cell sources including genetic irrelevance and limited supply^31,32^. Leveraging an isogenic iPSC cell line enabled us to systematically investigate and compare the host response across diverse regions of human lungs. Our results reveal a differential immune response between the human airway and alveolus against SARS-CoV-2. We also show that IAV causes multifold cellular damage and immune response as compared to SARS-CoV-2. By unraveling these complexities, our models offer a refined understanding of the innate immune responses associated with different viruses and disparate regions of the human lungs.

By culturing iPSC-derived CPM+ lung progenitor cells on microfluidic devices, we effectively obtained on-chip models for human airway and alveolus, each populated with distinct airway and alveolus cells, respectively. The airway epithelium consisted of multiciliated, mucus-secreting, and basal cells, while alveolus epithelium comprised of AT2 and AT1-like cells (Fig. 1b-k). This cellular composition closely mimics the *in vivo* cellular diversity of human lungs^43,44^. Co-culturing endothelial cells with respective tissues further enhanced the similarity to the *in vivo* interface of human airways and alveoli, as reported previous MPS studies^27–30^. Notably, the prevalence of multiciliated and AT2 cells in the respective chips underscores their efficacy in mimicking respiratory viral infections *in vivo*, given that these cell types are primary targets for a broad spectrum of respiratory viruses, including SARS-CoV-2 and IAV infection^38,45,46^.

The iPSC-derived epithelial cells in both chips exhibited permissibility to viral infections, including SARS-CoV-2, and IAV (Fig. 2c, d, f and 3a, e). Notably, previous MPS using primary cells failed to demonstrate productive SARS-CoV-2 infection on-chip, potentially owing to inadequate genotypic similarities to the *in vivo* characteristics of lung epithelial cells^27^. The consequential cellular damage observed in ciliated cells and AT2 cells in respective chips was markedly less pronounced post-SARS-CoV-2 compared to IAV infection (Fig. 2f, g and Fig. 4d, g). IAV infection resulted in a significant reduction in the cilia area covered and the GFP area covered in infected airway and alveolus epithelium, respectively. To the best of our knowledge, there are no reports within the MPS literature that have illustrated such comprehensive differences in cytopathies related to two distinct viral infections. This underscores the remarkable capability of our models to faithfully mimic cytopathic effects associated with different viral infections.

IAV infection induced damage the stratified layer of the airway epithelium, accompanied by an increased population of basal cells (Fig. 4d), suggesting a regenerative phase following substantial loss of multiciliated cells^47–49^. Co-localization of IAV nucleoprotein and cytokeratin further indicates the dedifferentiation of infected cells as reported previously^50^. In IAV-infected alveolus epithelium, individual cell sizes increased while maintaining the tight junctions (Fig. 4f). Notably, AT2 cells, pivotal in regenerating the alveolar space after lung injury, may undergo a transitional stage marked by an increase in cell size, termed hypertrophy^51–53^. The observed increase in cellular size may suggest a regenerative phase in the alveolus epithelium post IAV-induced acute injury. Significantly, the recapitulation of the post-infection regenerative phase in both airways and alveoli is reported for the first time in MPS models.

Capitalizing on the unique advantages of our isogenic induced pluripotent stem cells (iPSCs), we conducted a comparative analysis of immune responses across different regions of human lungs following SARS-CoV-2 and IAV infections. Our findings reveal a distinctive innate immune response triggered by SARS-CoV-2 in the human airways and alveolar tissue (Fig. 3b-f). Specifically, in airway chips, SARS-CoV-2 upregulates key genes and pathways related to type I interferon, whereas in alveolus chips, type I interferon was significantly suppressed. This dysregulated activation of type I IFN after SARS-CoV-2 infection aligns with previous reports^54^. Our system provides a valuable platform for comprehending the variations in immune responses between different regions of the human lung, an aspect that has not been well-understood^25^. The usage of iPSCs addresses irregularities related to SARS-CoV-2-related immune responses, such as non-productive viral replication and uncharacteristically high immune response reported in previous MPS reports^27,28^.

Remarkably, despite a suppressed type I response, chemokine pathways were still significantly upregulated in epithelium of SARS-CoV-2 infected alveolus chips (Fig. 3g). A delayed/suppressed IFN1 activation with upregulated chemokine pathways may lead to uncontrolled viral replication and recruitment of immune cells^34,55,56^.

The immune response elicited by both airway and alveolar epithelium to IAV infection was multifold higher than SARS-CoV-2. In IAV-infected airway and alveolus chips, there was a simultaneous upregulation of interferon pathways with widespread activation of antiviral and apoptotic pathways (Fig. 5c-g). The diminished immune response against SARS-CoV-2 is hypothesized to result from a more effective suppression of the innate immune response, particularly the type I interferon response, by SARS-CoV-2. Notably, the overall heightened immune response post-IAV infection correlated with a more extensive degree of cellular damage compared to SARS-CoV-2. The discernible differences in immune responses against different viruses underscore the capability of our isogenic induced pluripotent stem cell (iPSC)-derived lung models to faithfully emulate virus-specific responses.

In SARS-CoV-2-infected alveolus chips, we noted no discernible upregulation of antiviral pathways, possibly attributed to a lack of interferon (IFN) activation (Fig. 3d). The viral titer in alveolus chips post-SARS-CoV-2 infection remained consistently elevated until day 7, indicating a sustained peak of viral proliferation (Fig. 6b). Previous reports provide hypothesis that a dysregulated IFN activation can lead to uncontrolled viral growth in patients resulting in severe COVID-19^24^. Additionally, IFNB1 pretreatment of infected alveolus chips inhibited SARS-CoV-2 infection (Fig. 6c). IFNB1 as a therapeutics option was popular in the peak of COVID-19 pandemic owing to its ability to restrict viral replication^57^. Inhibition of viral infection by IFNB1 pretreatment not only substantiates the important role of type I interferon in viral inhibition but also provides a proof of concept of our model as a potential drug screening tool.

SARS-CoV-2 and IAV infection in endothelium were not observed in either the airway or alveolus chips at 3 dpi (Fig. 2c and Fig. 3a, e). RNA sequencing analysis of the endothelium in both SARS-CoV-2 infected chips indicated lack of immune response related genes upregulation (Fig. S4). While endothelial cells are rarely shown to be permissive to SARS-CoV-2 infection^58^, it has been reported that infected epithelial cells can release factors capable of eliciting immune response in the endothelial cells, especially after crossing the porous membrane that mimics the *in vivo* basement membrane^27,39^. The absence of immune response in endothelium could be attributed to the limited migration of cytokines from the top to the bottom channel, owing to a less prominent immune response in the epithelium. Conversely, RNA sequencing analysis of IAV-infected airway and alveolus chip demonstrated upregulation of genes and pathways related to type I interferon and other immune responses. (Fig. 6a). This aligns with prior reports of immune system activation in endothelial cells post-IAV infection of epithelial cells is reported previously^30^.

## Conclusion

Our iPSC-derived airway and alveolus chips closely replicate the *in vivo* interface of the human upper airway and alveolar sac. We achieved optimal differentiation of isogenic lung progenitor cells into airway and alveolar cells in respective chips. Our study reveals a tissue-specific response to IAV and SARS-CoV-2 infection.

We highlight a discernible difference of innate immune responses elicited by SARS-CoV-2 in our iPSC-derived airway and alveolus chip models, a phenomenon that is proved to be difficult to emulate using traditional immortalized or primary cell sources. Furthermore, we offer insights into the comparatively conservative nature of the immune response triggered by SARS-CoV-2 in contrast to IAV. With ongoing advancements in vaccine and drug administration methods, such as nasal delivery, our chips, derived from identical cell sources, hold tremendous potential for assessing their tissue-specific effectiveness in future, alongside the utility in therapeutic drug screening.

## Materials and Methods

### Microfluidic device fabrication

The microfluidic device consists of two Polydimethylsiloxane (PDMS) layers with microchannels measuring 1 mm × 1 mm, separated by a porous PET membrane (Falcon® Permeable 0.4 µm Transparent PET Membrane, 353090, Corning, USA). Soft lithographic technique was employed to make the PDMS layers with 1 mm × 1 mm channel. With 10:1 base to curing agent ratio, a PDMS prepolymer (Silpot 184; Dow Corning, USA) was cast against CNC milling machining fabricated mold.

To bond the two PDMS layers to the semipermeable PET membrane, liquid PDMS prepolymer was applied through spin-coating at 2500 rpm for 60 s directly on a glass slide. The thin layer of PDMS prepolymer was transferred to the PDMS layers with microchannels by placing them directly on the glass slide before sandwiching the membrane between the microchannels. The assembled PDMS layer with the PET membrane was then kept at 4 ℃ for 48 h to remove air bubbles, after which it was placed in an oven at 50 °C overnight to cure the PDMS prepolymer.

### iPSCs differentiation into NKX1+ lung progenitor cells

SFTPC-GFP knockin reporter iPSCs (B2-3)^36^ were differentiated into definitive endodermal cells in a stepwise manner, followed by anteriorization and ventralization to generate CPM+ lung progenitor cells in the specified medium in each step (Table S3), as previously described^59^. In brief, iPSCs were initially seeded on culture plates coated with Geltrex (Thermo Fisher Scientific, USA) in RPMI medium with 2% B-27 supplement (Thermo Fisher Scientific), containing 100 ng/ml Activin A (API Co., Ltd., Japan), 1 μM CHIR99021 (Axon 1386, Axon Medchem, Netherlands), 50 U/mL penicillin–streptomycin (Thermo Fisher Scientific), and 10 μM Y27632 (Y-5301, LC Laboratories, USA). On the following day, sodium butyrate (193-01522, Fujifilm Wako, Japan) was added to the medium to reach a final concentration of 0.25 mM. From Day 2 to Day 6, the cells were cultured in a medium supplemented with 0.125 mM sodium butyrate and without Y27632.

From Day 6 to Day 10 (Step 2), the cells were cultured in DMEM/F12 (Thermo Fisher Scientific) basal medium with 2% B-27 supplement, 1x GlutaMax (Thermo Fisher Scientific), 50 U/ml penicillin–streptomycin, 0.05 mg/mL L-ascorbic acid (Fujifilm Wako), and 0.4 mM monothioglycerol (Fujifilm Wako). During this period, the medium was supplemented with 100 ng/mL Noggin (R&D Systems) and 10 μM SB431542 (Fujifilm Wako) to induce anteriorization. From Day 10 to Day 14 (Step 3), the cells were cultured in the basal medium supplemented with 3 μM CHIR99021, 20 ng/mL BMP4 (Proteintech), and 0.05 μM all-trans retinoic acid (Sigma-Aldrich) to induce ventralization.

From Day 14 to Day 21 (Step 4), the cells were cultured in the basal medium supplemented with 3 μM CHIR99021, 10 ng/mL KGF (PeproTech), 10 ng/mL FGF10 (PeproTech), and 20 μM DAPT (Fujifilm Wako). Subsequently, the cells were labeled with a rat anti-CPM antibody (in-house) and microbead-conjugated anti-rat immunoglobulin kappa light chain antibody (130-047-401, Miltenyi Biotec). This was followed by the isolation of CPM+ lung progenitor cells using an autoMACS Pro Separator (Miltenyi Biotec) for the subsequent experiments.

### Differentiation on-chip

#### Airway-on-chip

Both channels of the devices were washed with 70% ethanol and dried at 50 ℃. Following UV treatment, both channels were rinsed with 200 μl of phosphate-buffered saline (PBS). Subsequently, the top side of the PET membrane was coated with 100 μl of iMatrix-511 (32 µg/ml) and incubated at 37 ℃ overnight. The following day, the coating material was removed by introducing 100 μl of PneumaCult™-ALI Medium (05001, Stemcell Technologies, Canada) without DAPT into the top channel, along with the addition of 300 μl in the bottom channel. The excess medium was removed from the top channel outlet reservoir using an aspirator.

Fresh CPM+ cells were suspended in PneumaCult™-ALI Medium without DAPT at a density of 2 × 10^7^ cells/ml and introduced into the top channel of each device. After 4 h, an additional 200 μl of PneumaCult™-ALI Medium without DAPT was directly added to the top channel. Once confluence was achieved, all the medium from the top channel was removed, and approximately 30 μl of PneumaCult™-ALI Medium with DAPT was introduced, along with 300 μl of PneumaCult™-ALI Medium with DAPT in the lower channel. The devices were then placed in an incubator at 37 ℃ for 28 days, with medium changes every other day.

The devices were maintained for at least 28 days before the lower channel was washed, and fibronectin (F4759, Sigma-Aldrich, USA) was introduced as a coating material for endothelial cells. Human umbilical vein endothelial cells (HUVEC) (cAP-0001, Angio Proteomie, USA) were suspended in EGM2 medium (CC-3162, Lonza, Switzerland) at a density of 1–2 × 10^7^ cells/ml. After 4 h, the PneumaCult™-ALI Medium with DAPT and EGM2 medium was added to their respective channels.

#### Alveolus-on-chip

The top side of the PET membrane was coated with 100 μl of Geltrex (Thermo Fisher Scientific) and left to incubate overnight. The following day, the coating material was removed by introducing 100 μl of alveolarization medium into the top channel, along with the addition of 300 μl in the bottom channel. Any excess medium was removed from the top channel outlet reservoir using an aspirator.

The alveolarization medium was prepared as follows: Ham’s F12 (Fujifilm Wako) supplemented with dexamethasone (50 nM), 3-Isobutyl-1-methylxanthine (IBMX) (100 μM), B27 supplement (1%), BSA (0.25%), HEPES (15 mM), CaCl2 (0.8 mM), ITS premix (0.1%), 8-BrcAMP (100 μM), human KGF (10 ng/ml), CHIR99021 (3 μM), SB431542 (10 μM), Y27632 (10 μM), and penicillin/streptomycin (50 U/ml) (Table S3). Frozen CPM+ cells, suspended in the alveolarization medium at a density of 2 × 10^7^ cells/ml, were introduced into the top channel of each device. After 4 h, alveolarization medium was added directly to the top channel. Once confluence was achieved, all the medium was removed from the top channel, and approximately 30 μl of fresh alveolarization medium was introduced. In the lower channel, 300 μl of alveolarization medium was replenished every alternate day. The devices were incubated at 37 ℃ for 5 days before the lower channel was washed, and fibronectin was introduced as a coating material for endothelial cells. HUVEC (cAP-0001, Angio Proteomie, USA) were suspended in EGM2 medium (CC-3162, Lonza, Switzerland) at a density of 1–2 × 10^7^ cells/ml. After 4 h, alveolarization medium and EGM2 medium were added to their respective channels.

### SARS-CoV-2 and influenza virus

The SARS-CoV-2 isolate (SARS-CoV-2/Hu/DP/Kng/19-027), which was supplied by the Kanagawa Prefectural Institute of Public Health, was cultured in VeroE6/TMPRSS2 cells. The viral titers were determined using a 50% tissue culture infectious dose (TCID_50_) method. The influenza A virus (A/California/04/09 [H1N1]) was propagated and titrated in MDCK cells through a plaque assay. All experiments involving SARS-CoV-2 were conducted within a biosafety level 3 containment laboratory at the Institute for Life and Medical Sciences at Kyoto University.

### Viral infection-on-chip

For virus infection on the chip, the top channel was carefully rinsed with PBS, and each device received 10^3^ TCID_50_ of SARS-CoV-2 or 10^3^ PFU of IAV in 30 µl of medium. Following a 1-h incubation at 37 °C, the virus solution was aspirated, and the top channel was washed twice with PBS. Subsequently, 200 µl of medium was introduced into the upper channel to sustain the epithelium. The medium in both the top and bottom channels was changed daily until the devices were either fixed for IFA or the cells were lysed for RNA examination.

### Viral titer experiment

Virus titers were examined using a TCID_50_ assay for SARS-CoV-2 and a plaque assay for IAV. Briefly, VeroE6/TMPRSS2 cells were prepared in a 96-well plate one day before conducting the assay. The virus samples were serially diluted and introduced into the cells, followed by an incubation at 37 °C for 4 days. The cells were monitored under a microscope to detect any cytopathic effects. TCID_50_/ml was calculated using the Reed–Muench method.

### RNA extraction from chip

For RNA extraction from the single-layer airway or alveolus chips, the top channel was washed with 200 µl DPBS. After washing, 350 µl cell lysis buffer (RNeasy Micro Kit (Cat. No. 74004; Qiagen)) was introduced in the top channel in two installments, and cell lysate solution was collected in a 1.5 ml tube. For bilayer chips, the top and bottom channels were washed with 200 µl each. After washing, 350 µl cell lysis buffer was introduced in the bottom channel in two installments, and cell lysate solution was collected in a 1.5 ml tube. Thereafter, a similar procedure was performed for the top channel and cell lysate was collected for RNA extraction and stored at -80 degree Celsius for further analysis.

### RT-PCR analysis

Total mRNA was isolated from the cell lysates using a RNeasy Micro Kit (Cat. No. 74004; Qiagen). The amount and quality of total RNA were assessed using a Thermo Fisher NanoDrop spectrophotometer. Subsequently, cDNA was generated from each sample employing the Prime Script RT Master Mix (Perfect Real Time) (Takara Bio Inc., Japan). Quantitative RT-PCR was performed using TB Green Premix Ex Taq II (Takara Bio Inc.) in an Applied biosystems Quantstudio 5, and β-actin (ACTB) was used as the housekeeping gene. The relative expression (fold change) of target genes was quantified using the double delta C method. The primer sequences used are provided in Table S2.

### Immunofluorescence microscopy

The top and bottom channels were washed with PBS before fixation with 4% PFA for 15 minutes. Following fixation, the cells were washed with PBS and permeabilized using 0.2% Triton X-100 for 15 min. Alternatively, devices were stored after fixation at 4 ℃ for a maximum of 7 days. A 5% goat serum blocking buffer was added to the permeabilized cells at room temperature after washing, and primary antibodies in the blocking buffer were introduced. The samples were stored overnight at 4 ℃ for 2 h at room temperature. A secondary antibody in blocking buffer was applied to the cells for 40–50 min after washing at room temperature. DAPI was used to counterstain the cellular nuclei after the secondary antibody step, and imaging was performed using confocal microscope (FV-3000; Olympus, Japan). Image processing was conducted using ImageJ. A list of antibodies used is provided in Table 1.

### RNA sequencing

The iPSC-derived airway and alveolus chips, with mock and virus infections (both SARS-CoV-2 and IAV), were utilized for RNA-seq analysis. Cell lysates were collected from both the top and bottom channels, as described earlier, using Buffer RLT (RNeasy Micro Kit (Cat. No. 74004; Qiagen). RNA extraction was performed from each cell lysate employing a miniprep kit (Qiagen). The RNA quality was analyzed using the 2100 Bioanalyzer (Agilent Technologies). Subsequently, each library was prepared following the manufacturer’s guidelines with the TruSeq Stranded mRNA Sample Prep Kit (Illumina). Sequencing was conducted using NextSeq2000(Illumina), and FASTQ files were generated via bcl2fastq-2.20. To prepare the data for analysis, adapter sequences and low-quality bases were trimmed from the raw reads utilizing Cutadapt ver. 4.1. The trimmed reads were then mapped to the human reference genome sequences (hg38) using STAR version 2.7.10a and a GENCODE (release 32, GRCh38.p13) GTF file.

Raw counts were calculated with htseq-count ver. 0.13.5, employing the GENCODE GTF file. Subsequently, a differential expression analysis was carried out using DESeq2 in R studio (R version 4.3.0). Differentially expressed SARS-CoV-2 and IAV genes were excluded in further analysis. Gene Ontology analysis was conducted using the Database for Annotation, Visualization, and Integrated Discovery (DAVID 2021)^60^. The generation of GO plots was achieved using the ggplot2 R package (version 3.4.2). Log_2_ fold change values for each gene were utilized to create a heatmap using GraphPad Prism (GraphPad software, USA).

### Image analysis and quantification

Cell number quantification: To quantify nuclei, we utilized 20× magnification DAPI channel images (636.396 μm × 636.396 μm) from at least 3 regions of interest (ROI) in 3 different devices. Z-stack images above the membrane level were employed for epithelial nucleus quantification with ImageJ. Initially, the z-stack images were imported into ImageJ and stacked. Subsequently, the image type was converted to 8-bit, and threshold was set. Watershed segmentation was performed to separate boundaries, and then categorized regions were counted using “Analyze Particles” function in ImageJ. The results were then plotted using GraphPad Prism (GraphPad software).

Cilia area percentage quantification: The area covered by multiciliated cells were measured by 60× magnification Alexa Fluor 488 channel images (Ac-Tub) (212.132 μm × 212.132 μm) from at least 3 ROI in 3 different devices. Cilia are specifically localized on the apical cell surface. Therefore, for analysis, the first 10 steps of z-stack images were considered. Initially, the files were imported into ImageJ and z-stacked. Subsequently, the file type was converted to 8-bit. The threshold was set, and the area was measured using built-in tools in ImageJ.

GFP area percentage quantification: To quantify GFP, we captured 60× magnification EGFP channel images (212.132 μm × 212.132 μm) from at least 3 ROI in 3 different devices. All images were adjusted for brightness using the same lookup table (LUT). Z-stack images were imported into ImageJ and stacked before being converted to 8-bit. The threshold was set, and the same procedure as for Ac-Tub was followed.

Layer thickness quantification: To determine the height of the cellular layer, 60× magnification DAPI channel images (212.132 μm × 212.132 μm) from at least three ROI in three different devices were used. The top z-stack image with DAPI fluorescence intensity and the last image before the intensity decline were manually selected. Images were acquired with a z-stack gap of 1 μm, so the total number of z-stacks was calculated to determine the layer height. The data was plotted on a graph using GraphPad Prism (GraphPad software).

### IFNB1 experiment

IFN-beta (300-02BC, Peprotech) was introduced into the top channel at a concentration of 10 ng/ml, 24 h prior to the infection experiment. Subsequent infection experiments were conducted following a similar protocol as described earlier.

### Statistics

A Student’s t-test was conducted using GraphPad Prism (GraphPad software) for RT-PCR data and image analysis (*p < 0.05, **p < 0.01, ***p < 0.001, ****p < 0.0001). For RNA-sequence data, further downstream analysis was carried out using criteria of P_adj_ < 0.05 and log_2_ fold change > 1. Unless otherwise specified in the figure legends and figures, a minimum of 3 devices were used for each statistical comparison.

## Acknowledgment

We thank our technical staff (Aki Kubo, Mayumi Morikawa, Megumi Kida, and S A Nyoman Putri Triantini) for fabricating the devices, cell cultures, and RTPCR experiments. We also thank Chiho Onishi for performing the infection experiments at the BSL3 facility. Dr. Takuya Yamamoto, Kazusa Ohkita, and Satoko Sakurai for RNA-seq library preparation and sequencing, and Dr. Koichi Igura and Koyori Ushitora for supporting the cell preparation.

## Author Contributions

Conceptualization: SY, KF, TN, SG, RY

Funding acquisition: TN, SG, RY

Methodology: SY, YM, TN, SG, RY

Resources: SY, TN, SG, RY, RM

Software: SY, KF

Investigation: SY, TT, YM, ST

Visualization: SY, KF

Supervision: TN, SG, RY

Writing-original draft: SY

Review and editing: SY, KF, TN, SG, RY

## Data and Material availability

Data supporting the findings of this study are available from the corresponding author upon reasonable request. The raw sequencing data files generated in the experiments can be found on NCBI Gene Expression Omnibus (GSE247677).

