## Supplementary Files for "Isogenic iPSC-derived proximal and distal lung-on-chip models: Tissue- and virus-specific immune responses in human lungs"

This file includes:

Supplementary Tables S1–S3

Supplementary Figs.S1–S4

**Table S1.** Antibody list

| Antibody | Manufacturer | Catalog No. | Concentration |
|---|---|---|---|
| Anti-Acetylated Tubulin | Sigma-Aldrich | T7451 | 1:4000 |
| Alexa Fluor® 555 Anti-Mucin 5AC antibody | Abcam | ab218714 | 1:200 |
| Alexa Fluor® Cyt5-647 Antibody | Abcam | ab193895 | 1:200 |
| Anti-ZO-1 | Thermofisher | 33-9100 | 1:300 |
| Anti-VE-Cadherin | Abcam | ab33168 | 1:300 |
| Anti-GFP | Aveslabs | GFP-1020 | 1:500 |
| Anti-CPM | Gotoh lab | 43A1 | 1:100 |
| Anti-SLC34A2 | A kind gift from Dr. Ritter Gerd | MX35 | 1:100 |
| Anti-HT1-56 | Terrace Biotech | TB29 AHT1-56 | 1:150 |
| Anti-IAV NP | Genetex | GTX125989 | 1:1000 |

**Table S2.** Primer sequences used for RT-PCR.

| Primer | Sequence |
|---|---|
| *SNTN* | GCTGCAAACCCAATTTAGGA<br>TGCTCATCAAGTTCAGAAAGGA |
| *FOXJ1* | CCTGTCGGCCATCTACAAGT<br>AGACAGGTTGTGGCGGATT |
| *SPDEF* | AAGTGCTCAAGGACATCGAGA<br>AGGAGCCACTTCTGCACATT |
| *SCGB1A1(CCSP)* | CACCATGAAACTCGCTGTCAC<br>AGTTCCATGGCAGCCTCATAAC |
| *CHGA* | CGGATCCTTTCCATTCTGAG<br>ACCGCTGTGTTTCTTCTGCT |
| *P63* | ACTGCCAAATTGCAAAGACA<br>TGACTAGGAGGGGCAATCTG |
| *NKX2-1(TTF-1)* | AGGACACCATGAGGAACAGC<br>GCCATGTTCTTGCTCACGTC |
| *ACTB(β-ACTIN)* | CAATGTGGCCGAGGACTTTG<br>CATTCTCCTTAGAGAGAAGTGG |
| *NRP1* | AACAACGGCTCGGACTGGAAG A<br>GGTAGATCCTGATGAATCGCGTG |
| *IFNB1* | CAT TAC CTG AAG GCC AAG GA<br>CAG CAT CTG CTG GTT GAA GA |
| *INFL1* | AACTGGGAAGGGCTGCCACATT<br>GGAAGACAGGAGAGCTGCAACT |
| *ISG15* | GCAGATCACCCAGAAGATCG<br>GGCCCTTGTTATTCCTCACC |
| *ISG56* | CCTTGCTGAAGTGTGGAGGA<br>CCAGGCGATAGGCAGAGA |
| *MxA* | CTTATCCGTTAGCCGTGGTG<br>CAAGGTGGAGCGATTCTGAG |
| *SFTPB* | GAGCCGATGACCTATGCCAAG<br>AGCAGCTTCAAGGGGAGGA |
| *SFTPC* | GCAAAGAGGTCCTGATGGAG<br>TGTTTCTGGCTCATGTGGAG |
| *PDPN* | TCCAGGAACCAGCGAAGAC<br>CGTGGACTGTGCTTTCTGA |
| *AGER* | GCCACTGGTGCTGAAGTGTA<br>TGGTCTCCTTTCCATTCCTG |
| *SLC34A2* | TCGCCACTGTCATCAAGAAG<br>CTCTGTACGATGAAGGTCATGC |
| *ACE2* | ACAGTCCACACTTGCCCAAAT<br>TGAGAGCACTGAAGACCCATT |

| | |
|---|---|
| *TMPRSS2* | GTCCCCACTGTCTACGAGGT<br>CAGACGACGGGGTTGGAAG |
| *N_Sarbeco*<br>(SARS-COV-2) | CACATTGGCACCCGCAATC<br>GAGGAACGAGAAGAGGCTTG |

28

**Table S3.** Composition of the culture medium used during different phases.

| Medium | Constituents | |
|---|---|---|
| Endodermitization medium (Step1) | Basal Medium | RPMI B27 supplement (2 %), Penicillin/Streptomycin (50 U/ml), Activin A (100 ng/ml) |
| | Chemical and cytokines | CHIR99021 (1.0 µM), Y27632 (Day 0; 10 µM), Sodium butyrate (NaB) (Day 1; 0.25 mM, Days 2 and 4; 0.125 mM) |
| Anteriorization medium (Step2) | Basal Medium | DMEM / F12, Glutamax, B27 supplement (2 %), L-ascorbic acid (0.05 mg/ml), Monothioglycerol (0.4 mM), Penicillin/Streptomycin (50 U/ml) |
| | Chemical and cytokines | Human Noggin (100 ng/ml), SB431542 (10 µM) |
| Ventralization medium (Step3) | Basal Medium | DMEM / F12, Glutamax, B27 supplement (2 %), L-ascorbic acid (0.05 mg/ml), Monothioglycerol (0.4 mM), Penicillin/Streptomycin (50 U/ml) |
| | Chemical and cytokines | ATRA 0.05 µM, human BMP4 (20 ng/ml), CHIR99021 (3.0 µM) |
| Step4 medium | Basal Medium | DMEM / F12, Glutamax, B27 supplement (2 %), L-ascorbic acid (0.05 mg/ml), Monothioglycerol (0.4 mM), Penicillin/Streptomycin (50 U/ |
| | Chemical and cytokines | CHIR99021 (3.0 µM), human FGF10 (10 ng/ml), human KGF (10 ng/ml), DAPT (20 µM) |

| On-chip culture | Airway chip | Alveolus chip |
|---|---|---|
| Basal medium | PneumaCult™-ALI Medium | Ham's F12, dexamethasone (50 nM), 3-Isobutyl-1-methylxanthine (IBMX) (100 µM), B27 supplement (1 %), BSA (0.25 %), HEPES (15 mM), CaCl2 (0.8 mM), ITS premix (0.1 %), 8-BrcAMP (100 µM) |
| Chemical and cytokines | Heparin (4 µg/ml)<br>Hydrocortisone (1.0 µM)<br>DAPT (10 µM)<br>Y-27632 (10 µM)<br>Penicillin/<br>Streptomycin (50 U/ml) | CHIR99021 (3 µM), SB431542 (10 µM), Y27632 (10 µM), human KGF (10 ng/ml), and penicillin/streptomycin (50 U/ml), |

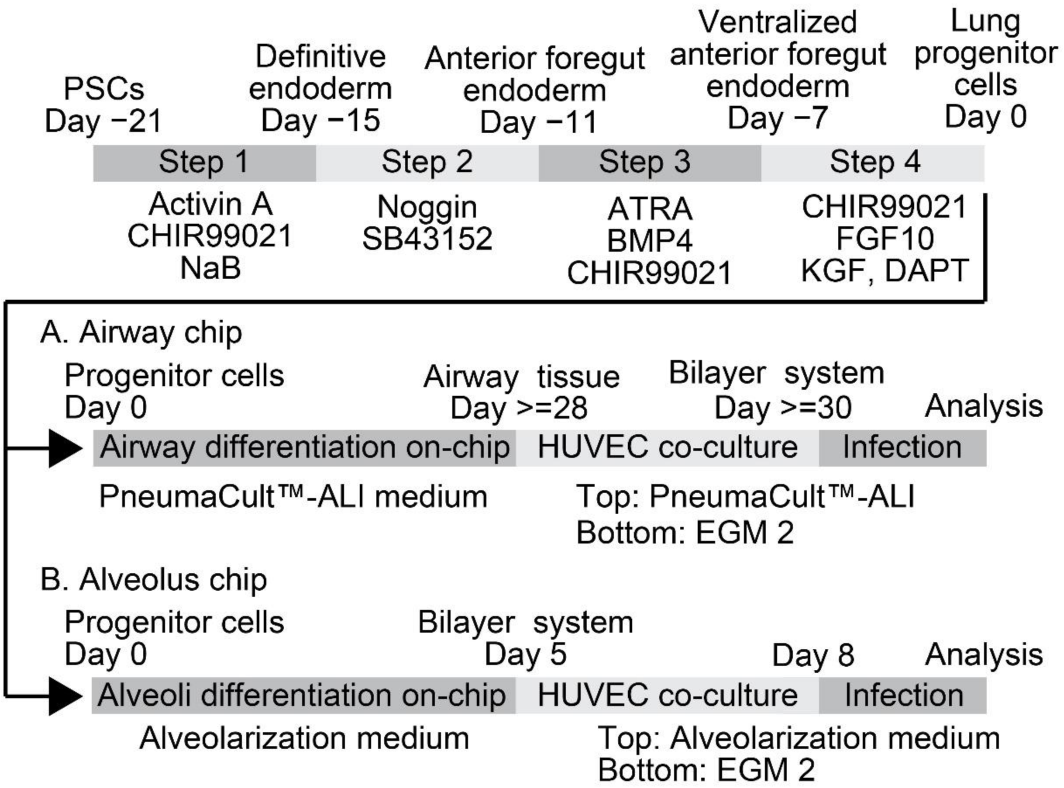


**Figure S1: Differentiation of iPSCs into airway and alveolar cells in microfluidic chips.**

Detailed protocol explaining the development of airways and alveolar chips using hiPSC-derived CPM+ lung progenitor cells.

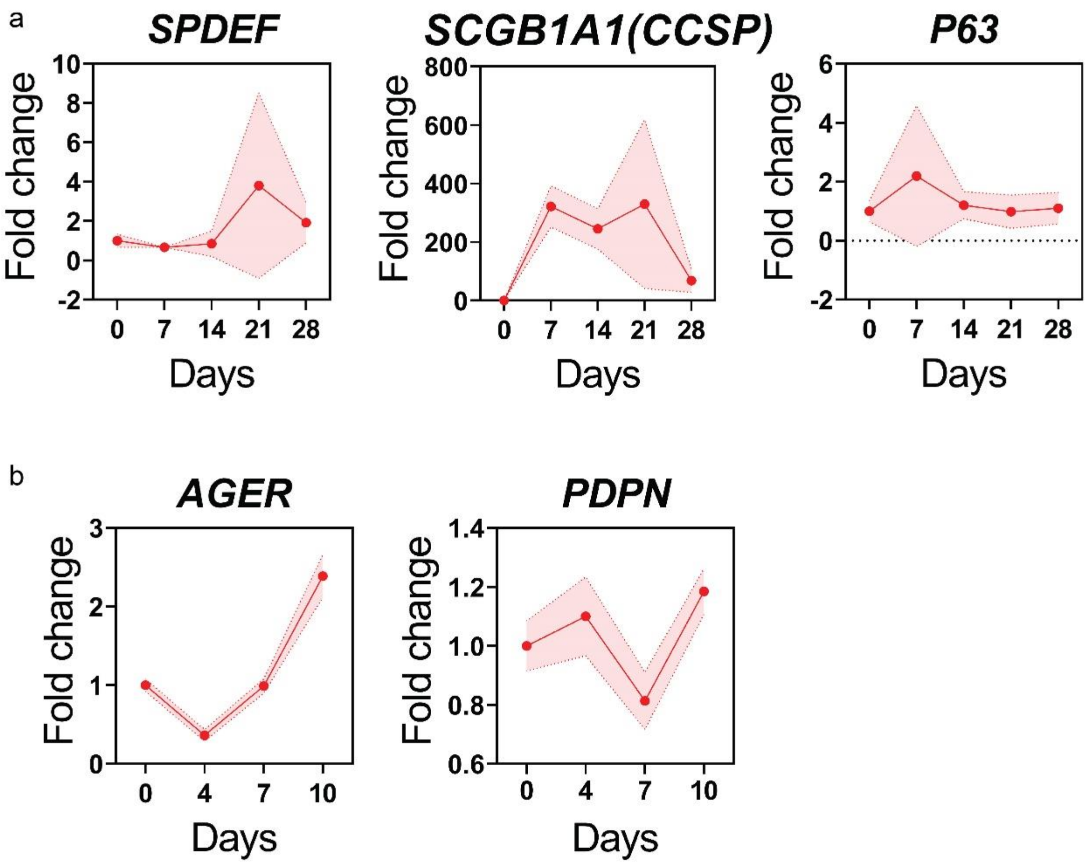


**Figure S2. Gene expression of mucus-secreting cells and basal cells in airway chips and AT1 cells in alveolus chip**

(a) Gene expression levels of *SCGB1A1(CCSP), P63,* and *SPDEF* in the airway epithelium and (b) *AGER* and *PDPN* in the alveolar epithelium (n = 3, except day 7, n = 2 in airway chips).

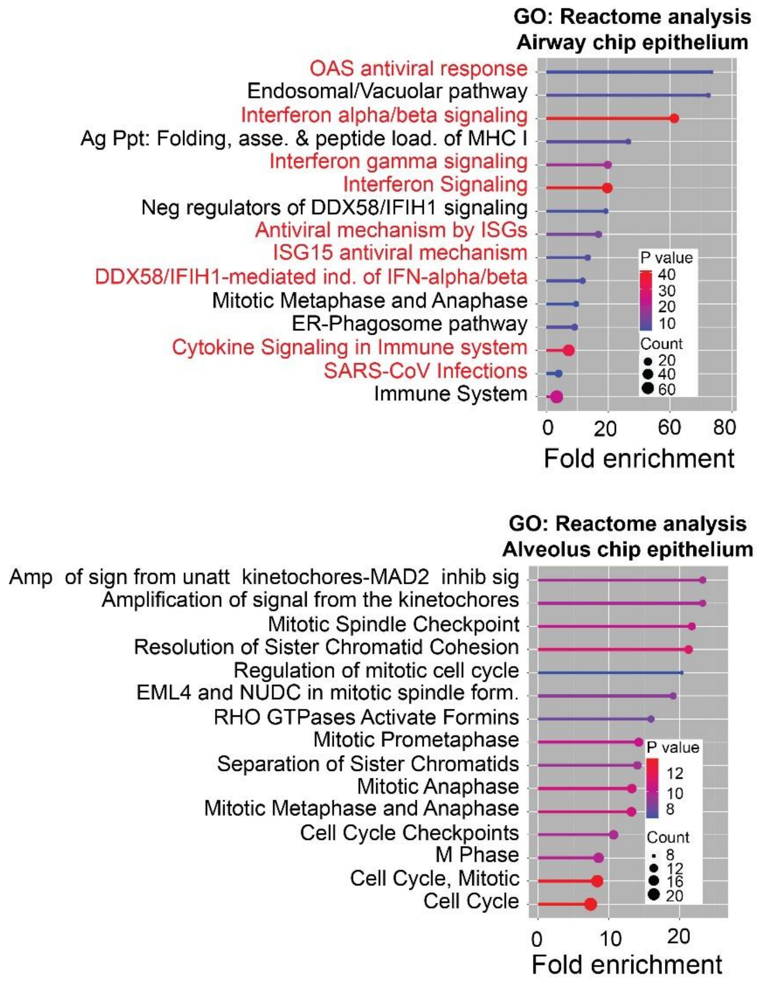


**Figure S3. IAV elicits innate immune response in both airway and alveolus chip epithelium.**

GO analysis revealed the most significantly upregulated reactome pathways in the airway and alveolus chips epithelium (P value shows $-\log_{10}$(P value)).

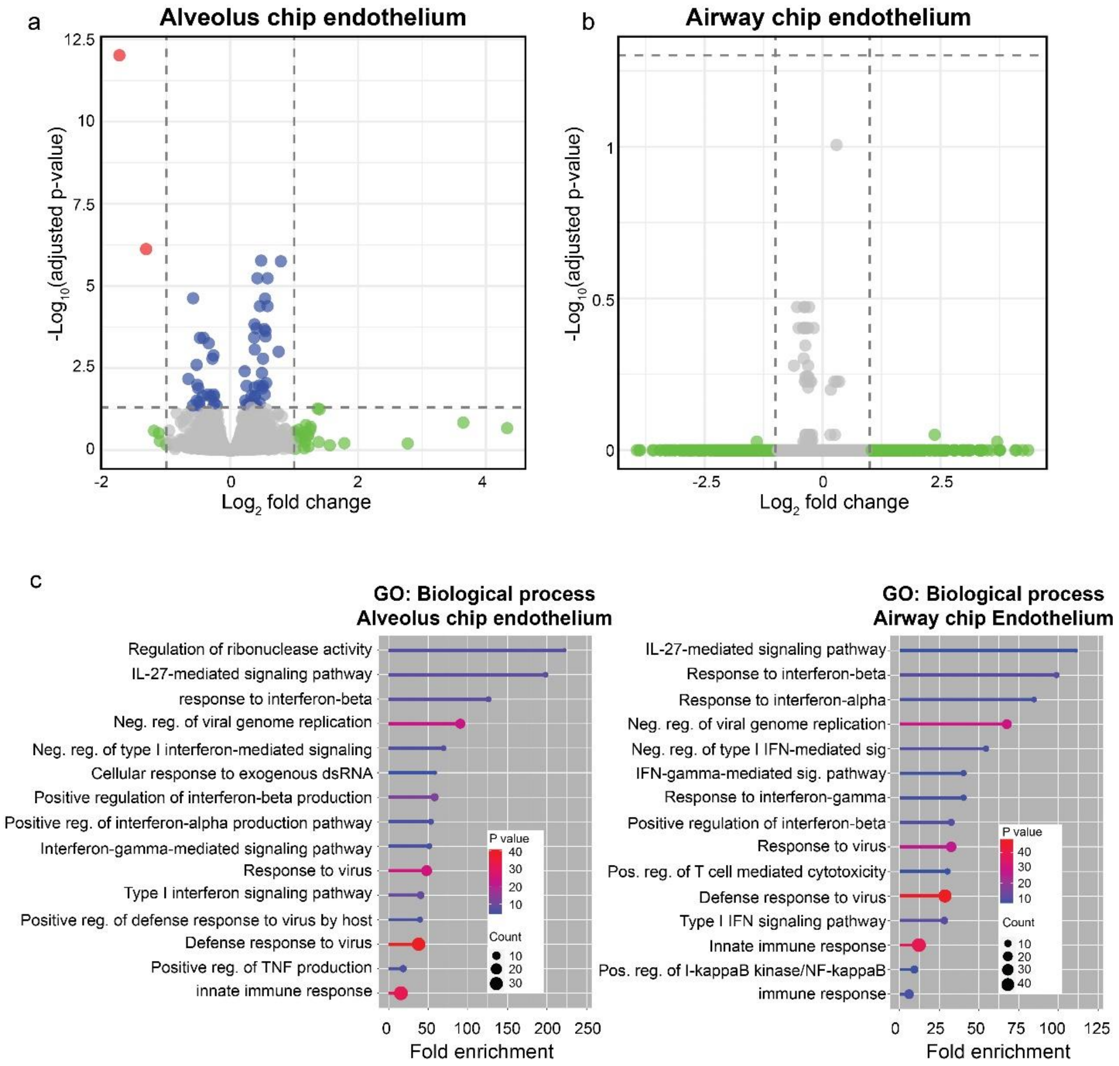


**Figure S4**. **IAV not SARS-CoV-2 infection induces relayed immune response in endothelial cells.**

(a) Volcano plot of differentially expressed genes between mock and SARS-CoV-2 infected alveolar endothelium ($\log_2$ fold change > 1 and ($P_{adj}$) < 0.05, n = 3 chips). (b) Volcano plot of differentially expressed genes between mock and infected airway endothelium ($\log_2$ fold change > 1 and ($P_{adj}$) < 0.05, n = 3 chips); (c) GO analysis displaying significant pathways upregulated in the endothelium of alveolus and airway chips between mock and influenza-infected samples (P value shows $-\log_{10}$(P value)).
